# Infection Tunes the Dynamics of Adenoviral E1A Disordered Regions

**DOI:** 10.64898/2026.04.12.717990

**Authors:** Patrick Koenig, Alexander Truong, Hadrien Lehman, Bailey-J C. Sanchez, Juris A. Grasis, Shahar Sukenik

## Abstract

Intrinsically Disordered Proteins and protein regions (IDPs) are abundant in many viral proteomes and play diverse roles in the viral infectious cycles. The adenovirus Early Protein 1A (E1A) is one such viral IDP. E1A acts as a molecular hub that regulates viral infection by mediating interactions between viral and multiple host proteins. Like other IDPs, E1A exists in a flexible ensemble of conformations. Despite a demonstrated link between ensemble structure and function in E1A, no real-time measurement of its ensemble has been performed. Here, we use live cell FRET microscopy to measure the local ensemble structure of E1A in human cells, both in healthy cells and in cells infected with adenovirus. We found specific disordered regions undergo significant changes to their ensemble in infected cells. Furthermore, infection also alters the propensity of these regions to partition between the cytoplasm and nucleus, a hallmark of E1A function during infection. Our results showcase that the structural ensembles of viral IDPs are responsive to infection, and suggest that these may play a role in regulating infection progression.

**Significance:** Integral to many viral proteomes are intrinsically disordered proteins and protein regions (IDPs), which target and rewire cellular pathways to ensure infection progression. During this process, the physical and chemical composition of the cell changes dramatically: metabolism is rewired, viral proteins are produced en masse, and as a result, the chemical composition of the host proteome is significantly altered. IDPs are known to be structurally sensitive to even mild changes in their environment, and their structural changes can result in a change to function. Here, using live cell FRET microscopy, we show that the structure and spatial localization of an adenovirus IDP, E1A, is altered in cells infected by the virus. Beyond a possible functional role for structural sensitivity in viral IDP function, our findings suggest that host IDPs may also be structurally altered by infection, with downstream functional consequences.

## Introduction

Viruses derive all resources required for replication and virion production by hijacking host cell pathways (1, 2). Remarkably, this hijacking uses an extremely limited number of viral proteins (often less than 30) (3). This means that few viral proteins must fill many functional roles, including the suppression of the host’s immune system, controlling its translation machinery, and rewiring of its metabolism to build and package virions (4, 5). Intrinsically disordered proteins (IDPs) are ideally suited to this task: a lack of stable three-dimensional structure results in rapidly interchanging conformations that are largely exposed to their surroundings. These expanded structural ensembles are interspersed with short linear motifs (SLiMs) that are designed to interact with host proteins (6), and can bind multiple partners with high fidelity (7, 8). Indeed, IDPs are commonly found in many viruses and are central to hijacking the host cell’s machinery (9).

Human Adenovirus type 5 encodes the Early protein 1A (E1A) - a viral IDP that acts as a molecular hub, mediating over 30 primary interactions and over 2,000 secondary interactions with host proteins (10). One of the most well-studied examples of this is E1A binding to cellular retinoblastoma protein – a key cell cycle regulator. E1A achieves this binding using two SLiMs that facilitate simultaneous and exceptionally tight binding (11–13). Key to this affinity is the accurate spacing of the two SLiMs, which is facilitated by a disordered region. Recent work showed that changes in the ensemble dimensions of this tether modulate the affinity of E1A to host retinoblastoma protein by orders of magnitude (14). This functional link between local ensemble dimensions and infection progression motivated us to study the local ensemble dimensions of E1A in the cellular environment.

Recent studies demonstrated IDPs are structurally sensitive towards perturbations in the cellular environment (15). This occurs because a lack of intramolecular bonds and extended surface area make solvent:protein interactions a dominant factor in determining ensemble conformations. Viruses introduce dramatic changes to the host cellular environment: Previous work has shown that during the timeline of infection, adenovirus causes intracellular acidification (16), ion and metabolite concentrations shifts (16–18), formation of noncellular compartments and condensates (19), and expression of nonendogenous proteins (20–22) (**Fig. S1**). Knowing that E1A function is affected by ensemble structure, and knowing that the changes in an infected cell’s physical-chemical composition can alter its ensemble structure, we wanted to understand if E1A ensembles are altered during the infectious cycle. More specifically, we hypothesize that in certain regions of E1A, the ensemble dimensions are altered over the course of the viral infection.

To test this hypothesis, we systematically probe the local structural ensembles of E1A in live, infected cells using genetically encoded reporters that rely on Förster Resonance Energy Transfer (FRET) to determine ensemble dimensions. We measure this FRET signal as cells are infected with human Adenovirus, tracking infected cells over a period of 48 hours. Our data shows that significant, persistent structural changes occur in specific regions of E1A upon adenovirus infection that do not occur in co-cultured, non-infected cells. In addition, we observed that structural changes are concomitant with shifts in nuclear localization, primarily driven by the C-terminal region of the protein. Taken together, our data shows that infection tunes E1A ensemble structure and localization, suggesting a previously unappreciated method of environmentally-driven regulation of viral IDP function. More broadly, our data points at the ability of IDPs to alter their conformation in response to distinct cellular states. Importantly, our findings prompt further investigations of how viral infection might also affect the structure and function of host IDPs.

## Results

### Tiling E1A systematically probes local ensemble structure of E1A

E1A is 289-residue long with an N-terminal Domain (NTD) and four conserved regions (CR) that are common across different adenoviral serotypes (23, 24) (**Fig. 1A**). Partitioned into each of these tiles are experimentally verified linear motifs that are critical to E1A versatility, as well as putative motifs that match known sequences, but may or may not be functional (**Fig. 1B** and **Table S1**). Prediction of E1A disorder using Metapredict v2.0 (25) showed a disorder score that is lower in regions containing known structures (helical regions in the NTD and zinc finger in CR3, **Fig. 1A**) (26, 27), while regions known to be disordered showed a high disorder score even in their tiled version (**Fig. 1C**).

**Figure 1.**
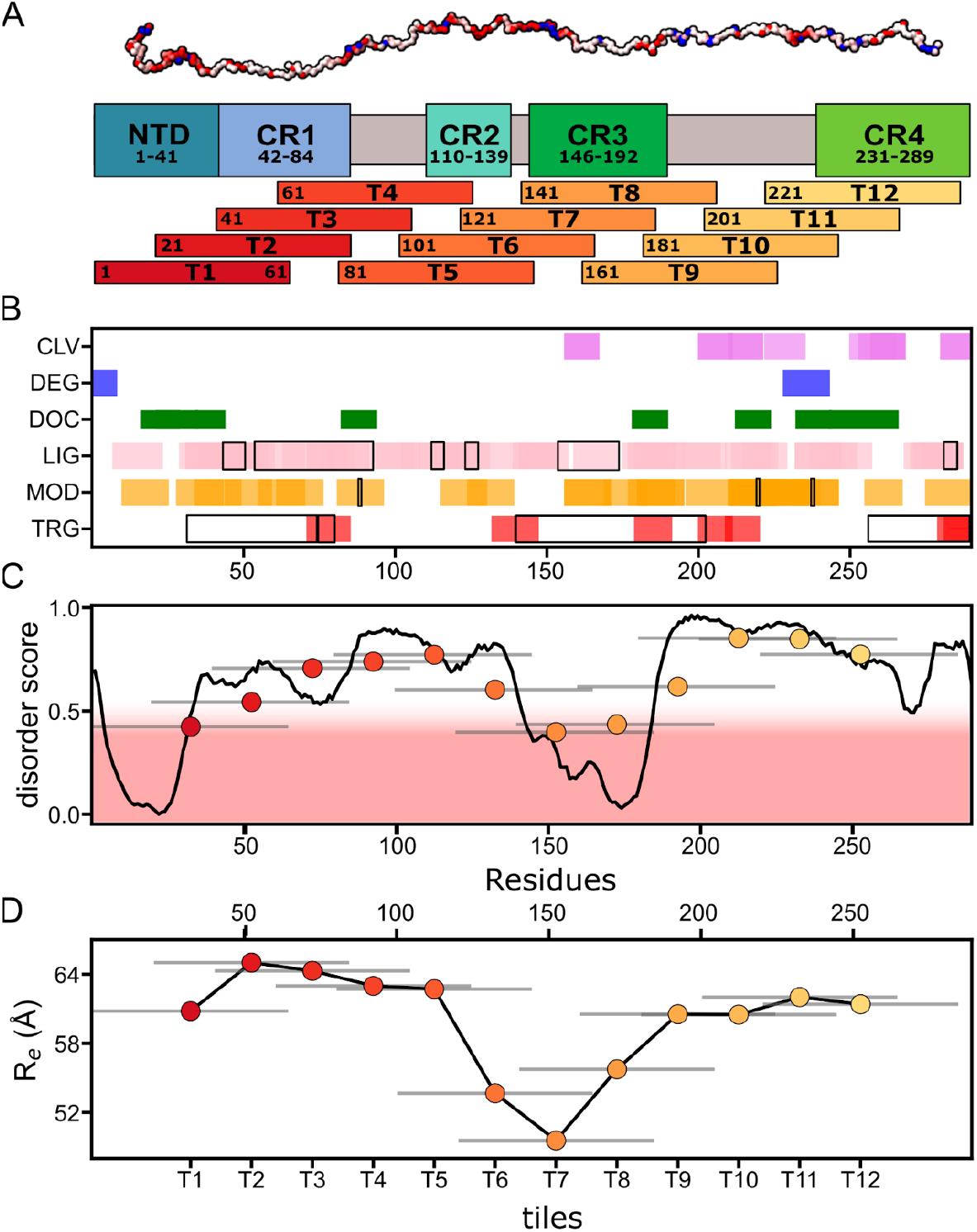
E1A tiles probe local sequence biases. **A**. E1A Domain map with tiling concept. (Top) Surface representation of the E1A backbone. Color gradient represents electrostatic charge: negative (red), neutral (white), and positive (blue) charged residues. (Middle) E1A domain structure, showing the N-terminal Domain (NTD) and Conserved Regions 1-4 (CR). (Bottom) E1A tiling. Each tile is 64-residues, with a 20-residue frameshift from its N-terminal neighbor. **B**. Linear motifs in E1A as found by the ELM database. Motifs include cleavage sites (CLV), degrons (DEG), docking motifs (DOC), ligand binding motifs (LIG), post-translational modification sites (MOD), and targeting sequences (TRG). Experimentally validated motifs are outlined in black and specified in **Table S1. C**. Disorder score calculated per residue using Metapredict v2.0 (black line). Scatter points are tile average disorder scores. Gray horizontal bars represent sequence coverage of individual tiles. Well-folded regions would fall in the red shaded regions. **D**. End-to-end distance (R_e_) predicted for each tile using ALBATROSS. Gray horizontal bars represent sequence coverage of individual tiles. Black line is a guide for the eye.

To understand how E1A ensembles change during infection, we opted for a tiling approach (tiles shown in yellow to red bars in **Fig. 1A**). Each tile consists of 64 amino acids, with a 40 amino acid overlap with the previous tile. This approach enabled us to focus on and evaluate the local conformational ensemble surrounding these modular linear motifs, as well as the local sequence chemistry in the cellular environment isolated from the full protein. We identified similar trends in the end-to-end distance, R_e_, using ALBATROSS (28) predictions for each of the tiles (**Fig. 1D**). Sequence chemistry evaluated by CIDER (29) further reveals a shift in tile charges (**Fig. S2A-B**) and charge distribution (**Fig. S2C**): a proliferation of negative charges at the N-terminal turns to a more positively charged C-terminal that can affect local structural biases within the cellular environment.

### E1A tiles display local structuring in the cellular environment

To quantify the ensemble dimensions in E1A tiles within the cellular environment, we used live cell FRET microscopy(15). To do this, we genetically encoded E1A tiles between two fluorescent protein genes: a donor (mTurquoise2 (30)) and an acceptor (mNeonGreen (31)). Once expressed, the fluorescent proteins form a FRET pair, and the ensemble dimensions of the tile connecting them determines the average distance between them. This distance is inversely proportional to the FRET efficiency, E_f_, calculated using the fluorescence intensity ratio of the acceptor to the total fluorescence intensity of both acceptor and donor (mTurquoise2 (32)) (**Fig. 2A**).

**Figure 2.**
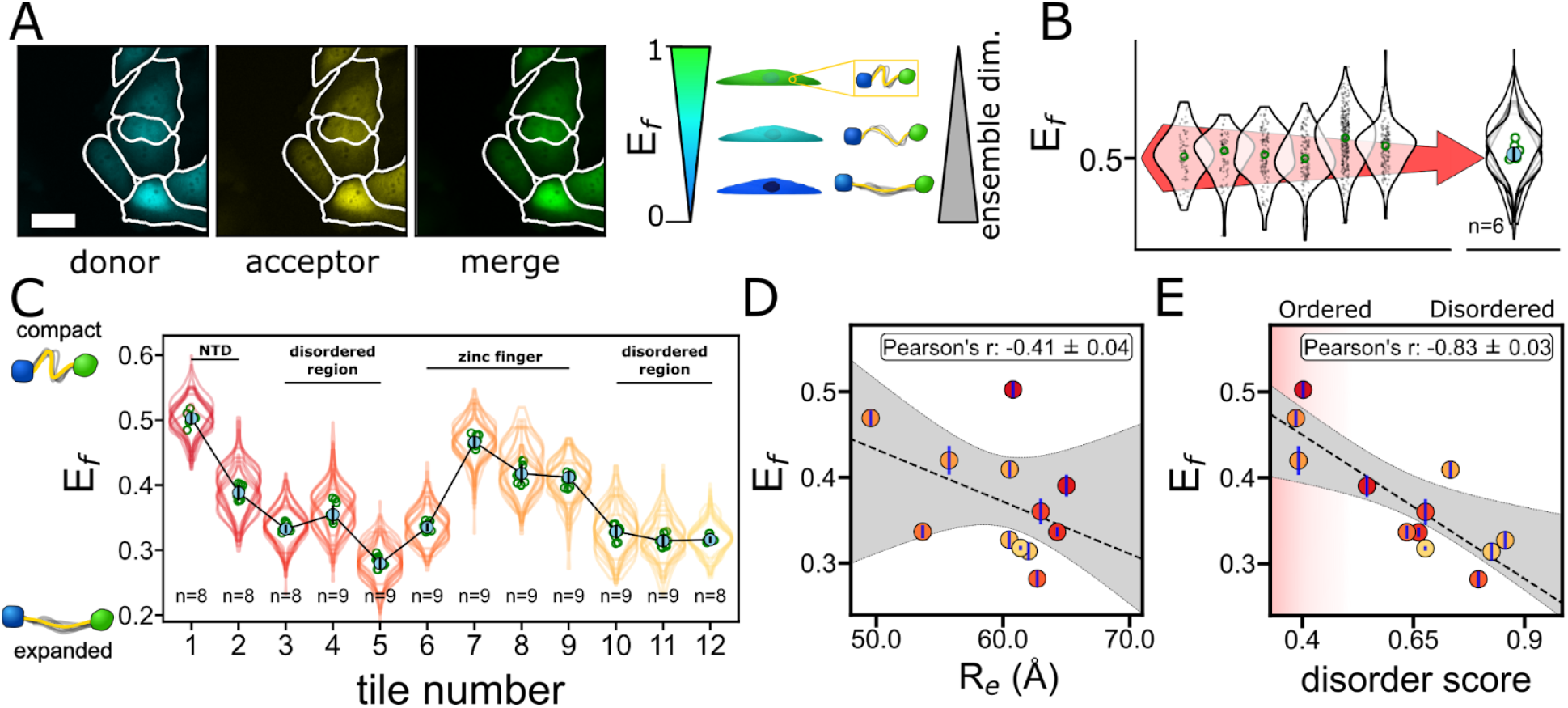
E1A tiles display local ensemble biases and localization changes. **A**. (Left) Microscopy images of cells expressing tile 1. Lines show cell segmentation by Cellpose. Scale bar is 30 µm. (Right) FRET Efficiency (E_f_) is inversely correlated with the IDR’s average end-to-end distance. **B**. Statistics used for violin plots. An individual biological repeat is a single imaging well. Within that well, imaged cells are segmented, and their average E_f_ is quantified (small grey dots) to create a violin plot (shown in the left panel) with the median shown as green circle. These violin plots are overlaid for the final representation of the tile (shown in the right panel), as well as the average of the medians of all individual violins (solid circle) and their standard deviation (black bar); n represents the number of wells imaged. **C**. FRET efficiency of E1A tiles. Features as in the overlaid violin in B. Trend line (black) links the average medians. **D, E**. Correlation between the measured FRET efficiency and end-to-end distance (R_e_, **D**) and Disorder Score (**E**) for each tile. Dashed line and shaded area represent a linear fit and 95% confidence intervals, respectively, and black bars are standard deviations of E_f_ averages shown in C.

The constructs were cloned into a lentiviral transgene plasmid under a CMV promoter and stably integrated into the cellular (U-2 OS) genome via lentiviral delivery. Once transformed, a population of cells expressing the construct was sorted to ensure uniform expression levels across all cells as measured by the fluorescence of the acceptor under direct excitation (DE). To quantify E_f_, fluorescence intensities for both donor and acceptor fluorophores are obtained by live cell imaging on a dual camera imaging setup, segmented using Cellpose (33), and averaged over the entire segmented cell area for donor and acceptor channels. To assess statistical differences between samples, we take a single imaging well in a 96-well imaging plate as a single biological repeat, reasoning that a well forms a distinct population for viral infection. All cells from each well are represented by a violin that shows the E_f_ distribution of all cell averages from that well. The median of each biological repeat is calculated, and the average and standard deviation of these medians are the reported values against which significance is assessed (**Fig. 2B**) (34).

We find that different tiles have a wide range of observable E_f_ values despite the same number of residues shared between all tiles (**Fig. 2C**). Tiles with high E_f_ are expected to be more compact, and this is in line both with low disorder prediction (e.g. NTD in tile 1) as well as with known helicity in the zinc finger (tiles 7-8). Further, we find that expansion and compaction trends persisted across multiple tiles. For example, tiles 2-5 and 7-10 showed increasingly expanded dimensions, while tiles 5-7 showed a compaction trend (**Fig. 2C**). Comparing these trends against known ordered regions, we see compaction increased when larger portions of the ordered region were encoded in the sequence, while greater expansion occurred in the disordered regions where SLiM occurrence is more frequent (**Fig. 1B,C**). We correlated tile E_f_ with the ALBATROSS predicted (28) end-to-end distance of the tiles, R_e_, (**Fig. 2D**) and found an expected negative correlation, with tile 1 showing outlying behavior. This is in line with other works showing partial structuring of the N-terminal domain (35). We further evaluated other sequence properties and found that compaction occurs in sequences where charged and proline residues are segregated (Ω, **Fig. S3**) (36). Finally, we found a direct correlation between E_f_ and predicted disorder (**Fig. 2E**). It’s important to mention that all sequence features and predictions are calculated in the absence of FPs. The correlations found between these and E_f_ imply that measurements are, at least to some degree, determined by the IDR properties rather than its artefactual interactions with the fluorescent proteins.

Together, these data indicates that the local ensemble structure of E1A, as predicted from the sequence and as reported in the literature from *in vitro* measurements, is recapitulated in the cellular environment: Tiles that were more compact (higher E_f_, tiles 1 and 7) had distinct sequence features or contained structural elements (35, 37). Conversely, tiles that were more expanded (lower E_f_, tiles 3-4, 10-12) coincided in regions reported to be disordered, where there was also a higher prevalence of SLiMs (**Fig. 1B**).

### E1A ensembles have local structural sensitivity to adenovirus infection

With basal dimensions of local E1A ensembles measured, we next wanted to measure if these ensembles change in cells infected with the human adenovirus serotype 5 (AdV5) virus. To ensure infection in imaged cells, we create a virus with a fluorescent reporter that would express in infected cells. To this end, we used the AdenoBuilder system (38) to exchange the AdV5 E3 gene region with mCherry, a red fluorescent protein (39) (**Fig. 3A**). Cells infected with this modified virus show a time-dependent increase in mCherry fluorescence intensity throughout the entire cell (**Fig. 3B, C**). We quantified the cell-average mCherry fluorescence levels and compared these to cells not treated with the virus (**Fig. 3D**) to discriminate between infected and non-infected cells (**Fig. 3E**). As a reference, cells that were not treated with virus established expected background mCherry fluorescence noise levels for non-infected cells at 12 hours (<300 AU, grey region in **Fig. 3D**).

**Figure 3.**
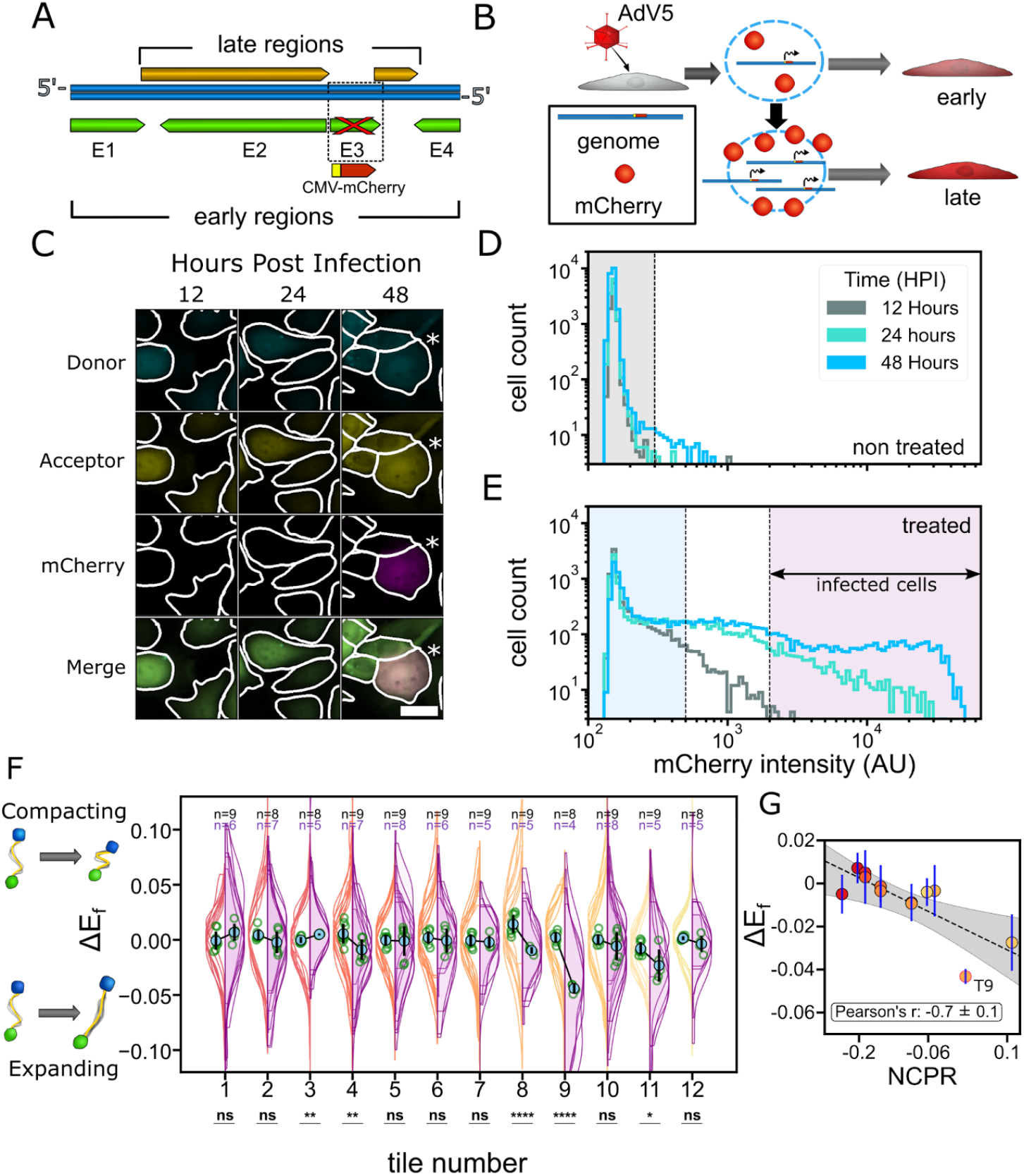
E1A tiles ensembles are structurally sensitive to infection. **A**. Diagram of adenovirus (AdV5) genome showing the replacement of the E3 gene region with a mCherry tag. **B**. Due to mCherry inclusion, cells infected with the modified AdV5 show a fluorescence intensity increase as infection progresses. **C**. Images of cells expressing a FRET construct that were infected by mCherry modified AdV5 over the course of 48 hours post-infection (HPI). Segmentation lines are shown in white. Scale bar is 30 µm. The cell labeled with an asterisk shows mCherry fluorescence due to AdV5 infection. **D,E**. Cell-average mCherry fluorescence intensity histograms for non-treated (D) and treated cells (E) at three time points post infection. Shaded areas denote mCherry fluorescence intensity cutoffs for non-treated (grey), non-infected (blue), and infected (purple) cells. **F**. Change in FRET efficiency between 48 and 12 HPI for non-treated (empty violins, left) and infected cells (purple violins, right). Asterisks denote significance as determined by a two-sided T-test. * p <0.05, ** p <0.01, *** p <0.001, **** p <0.0001. Violin features as in Fig. 2B. **G**. Change in FRET efficiency between 48 and 12 HPI for infected cells correlated with the net charge per residue for each tile. Markers are averages from the overlaid violins in (F), and error bars are the standard deviation. Dashed line and shaded area represent a linear fit and 95% confidence intervals, respectively.

In cells where the virus was introduced, we observed population shifts towards increased red fluorescence levels over the course of 48 hours. Over this time course, we measured cell-average mCherry fluorescence in both wells where no virus was added (non-treated), as well as where virus was added. In cells where no virus was added to the well, no change in fluorescence was observed over the course of 48 hours (**Fig. 3D**). Conversely, in wells where the virus was added some cells showed 1-2 orders of magnitude increase of mCherry fluorescence, indicating infection (**Fig. 3E**, purple region). Even in these wells, a population with fluorescent levels equivalent to non-treated cells was seen after 48 hours (**Fig. 3E**, blue region). We therefore labeled cells as “infected” if after 48 hours they displayed a high mCherry fluorescence (>2,000 AU, purple regions in **Fig. 3E**), and non-infected (<500 AU, blue region in **Fig. 3E**).

With our infection reporter system in place, we next compared E_f_ changes between infected and non-treated cells for each of our E1A tiles. We report on a change in FRET efficiency, ΔEf = E_f_ (48 HPI) - E_f_ (12 HPI), looking at infected cells (purple region, **Fig. 3E**) and non-treated cells (grey region, **Fig. 3D**). Our point of comparison was selected to be 12 HPI since no significant mCherry fluorescence was seen at this point, and allowed the closest meaningful comparison between infected and non-infected cells, preventing artifacts arising from depletion of non-infected populations at later timepoints. As the infected population begins to appear, we see distinct compaction and expansion of specific tiles compared to non-treated populations (**Fig. 3F**). The most distinguishable tiles are 9 and 11, which display a significant and strong expansion. We also point out that most other tiles that displayed significant changes, the ensembles seemed to compact rather than expand, though the magnitude of this change was small.

To try and understand the reason for the structural sensitivity of tiles to infection, we compared ΔEf with different sequence features across all tiles. We find little correlation between this sensitivity and the basal dimensions as measured in noninfected cells (**Fig. S4**). We also find little correlation with other sequence chemistry features, including average charge and charge distributions (**Fig. S5**) as well as lower correlations with other ensemble predictions and chemistry (**Fig. S5**). We do find, however, that both tile 9 and 11 have a low but positive net charge per residue, whereas all other tiles have a negative net charge per residue (**Fig. 3G**), and as a result an outlying isoelectric point (pI in **Fig. S5**). Compared to a pI of 3-4 for all tiles, Tile 9 with a pI of 7.8 seems especially poised to sensitivity when the surrounding pH changes. Our data suggests that the large change in dimensions could be a result of intracellular pH change, which has been previously reported to occur in the intermediate stages of adenovirus infection (40).

### Nuclear localization is non-localized and varies along E1A sequence

Localization of E1A to the nucleus is required for efficient viral DNA replication (41–43). It is well known that IDPs can contain multiple localization signals within a single protein, and that the balance between these localization motifs determines the overall spatial distribution of the protein across the cell (44–46). Previous studies indicate that E1A contains multiple nuclear inclusion and exclusion signals (NLS and NES, respectively). This includes a C-terminal NLS region with two potential weaker NLS in the N-terminal region and CR3 (47, 48), and also one validated NES in the CR1 between residues 70-80 (**Fig. 4A**) (49). Having shown that regions of E1A change the structure during infection, we next wanted to see if this also occurs for the partitioning of these regions between nucleus and cytoplasm.

**Figure 4.**
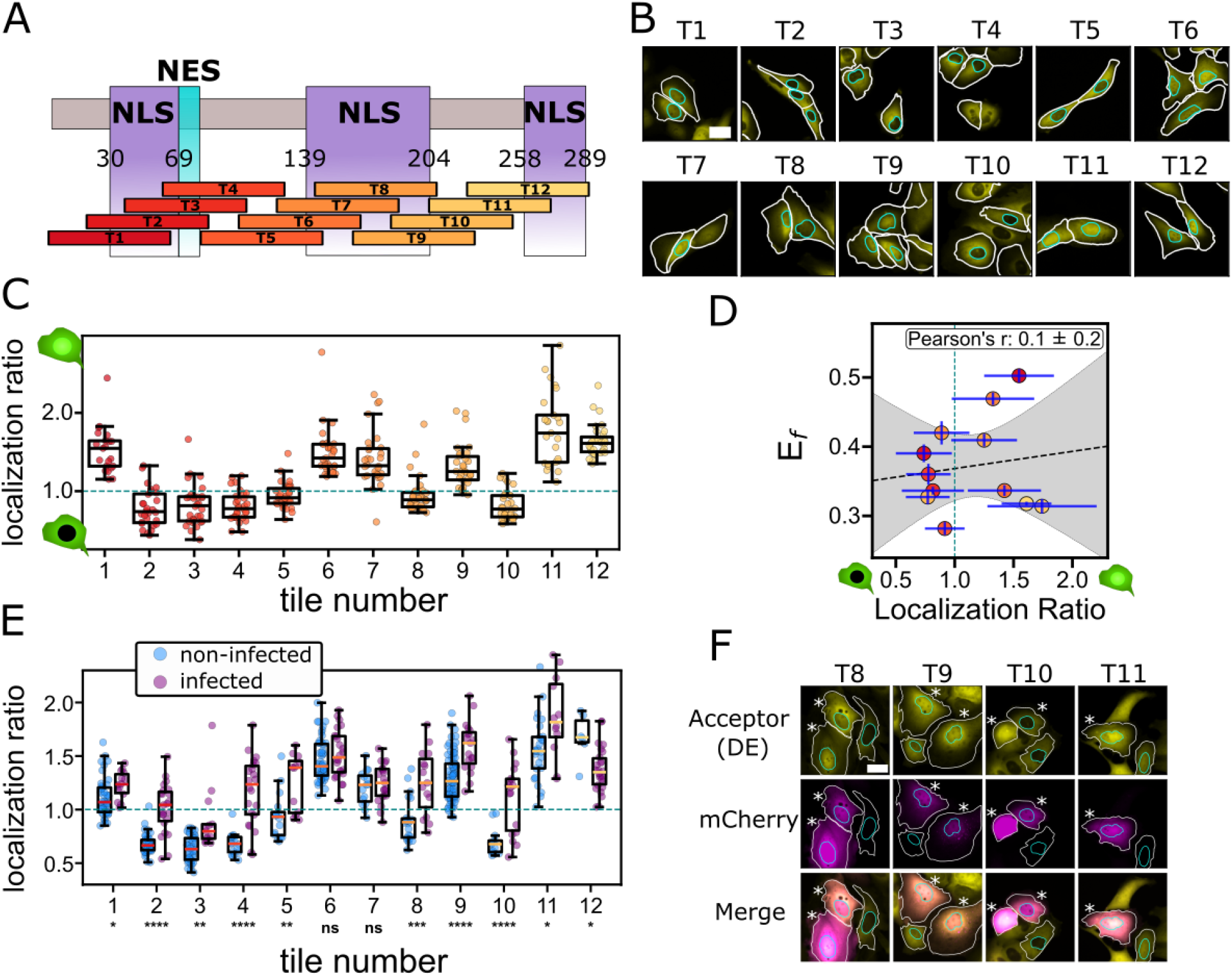
Nuclear localization changes during late-stage infection. **A**. E1A experimentally validated nuclear localization (NLS, purple) and exclusion (NES, turquoise) regions, and the E1A tiles in which they are included. **B**. Non-infected cells under direct acceptor excitation (DE). Line shows segmentation of whole cell (white) and nucleus (cyan). Scale bar is 30 μm. **C**. Ratio between the fluorescence in the nucleus and cytoplasm for each tile, taken from direct acceptor illumination. Box represents median 50% of the data, and line represents the median. Error bars span 90% of the data, and circles are individual cells. Dashed horizontal line indicates even DE fluorescence in cell and nucleus. **D**. Correlation between E_F_ and average localization ratio for each tile. Dashed line and shaded regions are linear fit and 95% CI of the data. **E**. Comparison of localization ratio in noninfected (blue, <500 AU) and infected populations (purple, >1500 AU). Asterisks denote significance between the populations, tested by an independent two-sided t-test. *p <0.05, **p <0.01, ***p <0.001, ****p <0.0001. **F**. Segmentation of infected and noninfected cells at 48 hours. Cell (white), nucleus (Cyan), Scale bar (30 μm). * denotes infected cells.

To do this, we looked at tile localization by measuring the acceptor emission under direct acceptor excitation (DE) in both the nucleus and the cell for each tile. It was visually apparent that there are tile-dependent differences in this localization, with some sequences showing very low fluorescence intensity in the nucleus (**Fig. 4B**). To quantify this, we segmented regions in the nucleus and cytoplasm of individual cells and took the intensity ratio between their average DE fluorescence. Our imaging revealed distinct localization patterns that differed between the tiles (**Fig. 4C**). An initial nuclear localization in tile 1 turned into exclusion in tiles 2-5 (residues 21-144). Localization then occurs again between tiles 6 (101-164) and 7 (121-184), encompassing residues 101-184. Unexpected exclusion occurred in tiles 8 (141-204) and 10 (181-244), while overlapping tiles 9 (161-224) and 11 (201-264) were found to localize.

To examine the determinants underlying localization differences between tiles, we looked for correlations between the localization ratio and different molecular features (**Fig. S6**). However, no clear correlations with sequence features were found. In addition, we found that localization is poorly correlated with ensemble dimensions (**Fig. 4D**). On the other hand, we do find that the existence of previously reported NLS and NES signals (**Fig. 4A**) effectively modulates localization: The NES signal is included in tiles 2-4 and effectively shuttles these tiles out of the nucleus. The N-terminal NLS is enough to provide localization in the absence of the flanking NES for tile 1, but is then insufficient to alter localization when the NES is included. The C-terminal NLS drives strong inclusion in tiles 11 and 12, and the central NLS correlates with localization in tiles 6 and 7, but surprisingly not for tile 8 which spans its entire length. Tile 9, which showed extensive structural change upon infection (**Fig. 3F**) shows significant nuclear localization despite being flanked by two tiles that are largely excluded from the nucleus. Overall, our data shows that subcellular localization is strongly correlated with previously reported localization signals, but these are difficult to predict from sequence or ensemble features alone.

We next wanted to see if localization might change in response to infection. To quantify localization changes, we looked at DE nuclear/cytoplasmic ratio in individual cells taken from infected cells (purple region in **Fig. 3E**) and non-infected cells (light blue region in **Fig. 3E**). From this comparison (**Fig. 4E, F**), we observed that in most tiles nuclear localization increases significantly. This change in localization is not correlated to ensemble dimensions (**Fig. S7A**). However, it is noticeably more pronounced in tiles that were excluded from the nucleus in non-infected cells (where the ratio is smaller than 1) (**Fig. S7B**). We further evaluated this change in localization to the various sequence properties (**Fig. S8**) and, again, found no clear strong correlation for infection-related accumulation. Overall, no clear correlation was observed for this infection-induced increase in nuclear localization.

Taken together, our results indicate that different regions of E1A have different tendencies to localize into the nucleus. Previously found NLS and NES, identified via deletions in the full-length protein, seem to have a significant effect on this localization. In addition, while nuclear localization increases over the course of 48 hours for most tiles, the extent of this localization is not uniform, and the underlying properties driving this change are not clear.

## Discussion

In this study, we investigated structural and spatial modulation of the Adenoviral E1A protein to better understand viral IDP dynamics in the infected host cellular environment. Local structural tuning that co-occured with infection was identified across the E1A sequence in multiple regions, in particular residues 161-224 close to the C-terminus (tile 9). This region demonstrated significant expansion that did not occur in any other tile of the sequence. Furthermore, certain regions displayed different tendencies to preferentially localize in the nucleus following infection. These were apparent in residues 61-100 and in the same C-terminal region where structural changes occurred in residues 161-224.

Examination of tile 9 sequence (**Table S2**) reveals that this tile has a region of net negative charge that is surrounded by positive charges that result in an overall slightly positive net charge, and a pI of 7.8 - unique features in the otherwise uniformly negatively charged E1A sequence (**Fig. S2A**). It has been reported in the past that tracks of positive charges can help localize regions into the nucleus (50). Furthermore, it’s been recently reported that tracks of charged amino acids may display non-cannonical charged states, modulating their overall charge (51, 52). We propose that previously reported changes that occur in the physical-chemical composition of cells infected with adenovirus may alter these charged states, leading to a change in conformation (40) (**Fig. 3E**). In line with this, past work has highlighted 6 dipeptide Glu-Pro repeats as essential to binding host transcriptional machinery (53, 54). Such tracks of negative charges may display a variation in charged states as the host cell pH changes from infection-induced metabolic rewiring (55). Any changes in the balance of charges in this region have been shown to affect its transcriptional regulation function (53) -providing a possible mechanism for both structural and functional changes.

That being said, other mechanisms explaining the ensemble changes we observe in infected cells should be considered. The SLiMs occurring in independent tiles already showed an agreement with their localization; it is therefore possible to consider other functionalities, including post-translational modifications, binding to cellular or viral components, or changes in *E*_*f*_ signal as a result of protein cleavage may be driving some of our observed changes. Indeed, multiple instances of putative (**Fig. 1B**) and experimentally detected linear motifs (**Table S1**) exist along the E1A sequence (**Fig. 1B**). Future mutational studies can be designed to decouple the impacts of local structural changes during infection from that of changes in linear motifs.

E1A acts primarily as a host transcriptional regulator: N-terminal SLiMs and motifs target pathways that result in cell cycle disruption and infection progression (18, 56, 57). Concomitantly, C-terminal features counteract this by driving cell differentiation pathways that delay infection progression (57, 58). Recent research has shown that altering ensemble dimensions, especially in transcriptional regulators activation domains, can lead to functional changes (59, 60). We propose that the ensemble changes observed in the C-terminal region of E1A help regulate the balance of functions that are required for viral progression. This can be accomplished, for example, by increased exposure of IDR regions that enhance SLiM accessibility.

Importantly, transcription regulation can only happen within the nucleus, making nuclear localization an additional factor affecting viral interference with host transcriptional machinery. We show that nuclear inclusion increases in infected cells for nearly all tiles of the protein, with specific regional increases in residues 141-264 (tiles 8-11) that relate to the previously described CR3 NLS and a portion of the bipartite C-terminal NLS (47, 61). This increase also coincides around key residues 202-208 that have been suggested to play a role in E1A localization (62). Outside the conventional C-terminal NLS, other localization signals have not been extensively characterized.

It is important to note some limitations in this study. First, by focusing on tiling the full-length protein, we lose the ability to observe any long-range interactions that could alter the behavior of the protein. Especially for localization, it is the interplay between multiple localization and exclusion signals that ultimately dictates the subcellular distribution of the protein. However, it is important to point out a large body of work showcases the modularity of IDPs, including direct measurement of tile function in hundreds of transcription factors (63–65) as well as in E1A (14), but it would be important to study the features found here in the context of the full-length protein to test their impact. An additional limitation is the attachment of fluorescent protein (FP) probes which may introduce artifacts into our FRET ensemble measurements and potential localization. While we cannot rule out intramolecular interactions between the IDP and FPs (and indeed these are likely to occur), a significant correlation between the predicted R_e_ (**Fig. 2D**) and the disorder score (**Fig. 2E**), predicted in the absence of FPs, indicates that, at the very least, the ensemble dimensions are not primarily defined by these interactions. Further, localization, though not directly implicated in ensemble-based exclusion (**Fig. 4D**), may impact the exposure of NLS in tiles 8 and 10 that would need to be verified in future studies.

Our findings point to the possible changes in ensemble:function relationship of IDPs in a cellular environment altered by viral infection. It also suggests these changes may act as a previously unknown mechanism regulating viral infection. Many clinically significant viruses, including coronavirus (66), herpes simplex virus (67), and human papillomavirus (68), possess these structurally disordered elements that carry pivotal roles in their life cycle. Perhaps most importantly, over 70% of human proteins contain IDPs (69), and their presence is especially over-represented in transcriptional regulators (70, 71). Like viral IDPs, host IDPs may also be altered in infected cells. Unlike their viral counterparts, host IDPs are not designed to function in an infected environment. Where such ensemble changes occur in the infected host’s proteome, dysfunction may follow. Understanding how IDP ensembles function in these altered environments may reveal new modes of host protein dysfunction during viral infection.

## Data, Material, and Software availability

All spreadsheet data, scripts, and software are available in the github repository (https://github.com/sukeniklab/Koenig_et_al).

## Acknowledgements

This research was initially funded through a University of California, Merced, Health Sciences Research Institute seed grant. Additional funding was provided by the National Institute of Health under award R35GM137926 to S.S., and by the National Science Foundation BII:2119968 to J.G., S.S. acknowledges support from the Sloan Foundation. P.K. was supported by a fellowship from NSF-CREST CCBM at UC Merced. We thank Dr. F. Bunz for donating the plasmids and assisting in the implementation of the Adenobuilder system. We acknowledge the Stem Cell Instrument Foundry at UC Merced and Dr. D. Gravano for use of the Aria III flow cytometer in cell sorting.

## Author Contributions

P.K., J.G., and S.S. designed research, P.K., A.T, H.L., and B.S. performed research, P.K. and S.S analyzed the data, and P.K., J.G., and S.S. wrote the paper.

## Competing Interest

The authors declared no competing interests.

## Notes

https://github.com/sukeniklab/Koenig_et_al

### Competing Interest Statement

The authors have declared no competing interest.

