## Supplementary figures and tables for "Infection Tunes the Dynamics of Adenoviral E1A Disordered Regions"

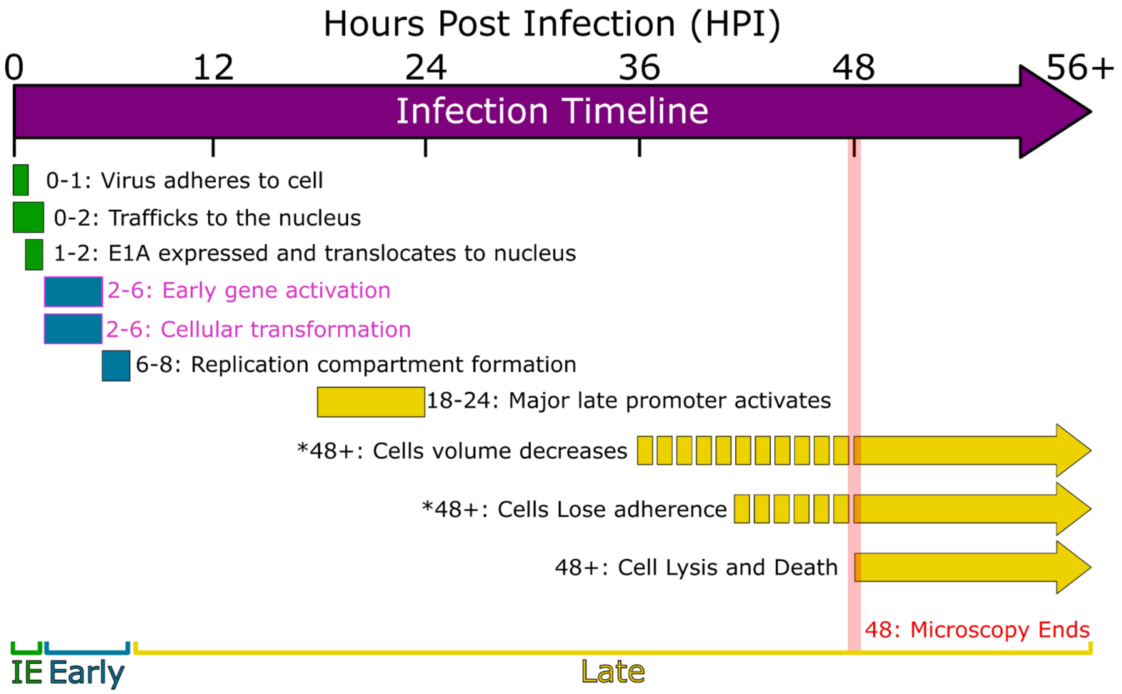


**Figure S1. Adenoviral intracellular infection timeline.** Key viral stages broken down by viral lifecycle, Immediate early (IE, green), early (blue), late (yellow) (1–3). IE to Early transition is based on early gene activation, Early to Late is dependent on DNA synthesis initiation. Solid bars represent the range of time at which events are likely to occur. * denotes observable events during microscopy in U-2 OS cells; broken bars represent low frequency of observation occurring in those time periods. Late events represented by an arrow indicate high variability of event occurrence. Events driven by E1A during the early life cycle are highlighted (pink).

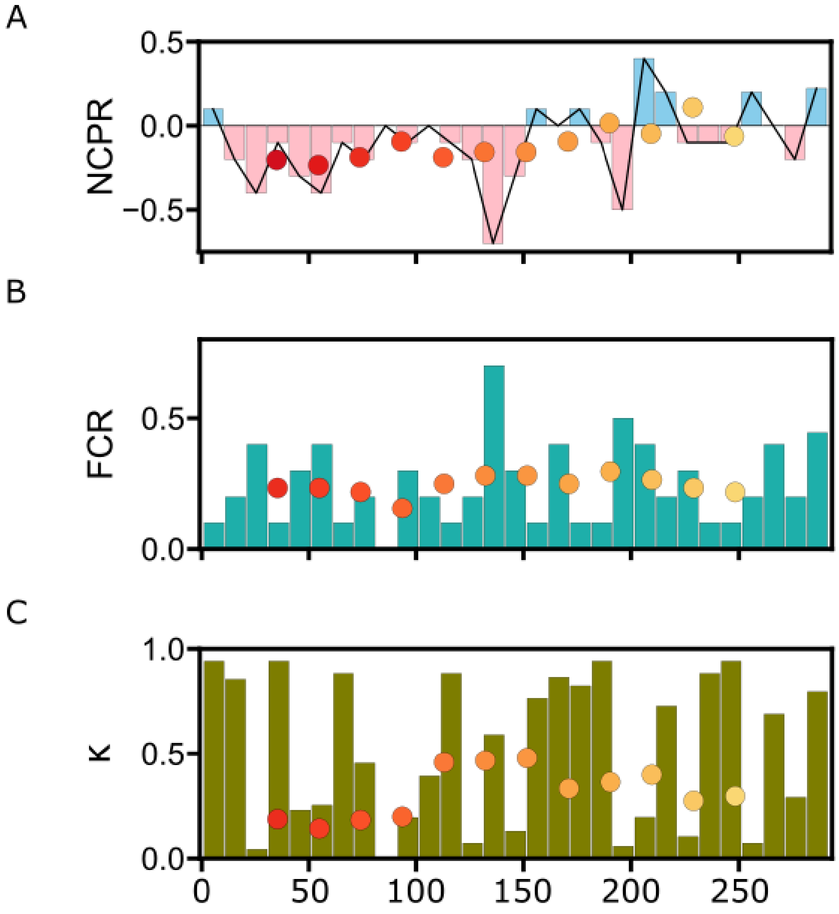


**Figure S2. E1A sequence chemistry analysis.** A-C. Bars represent the 10 amino acid average, overlaid scatter points are E1A tiles. A. Net charge per residue. Trend line (black). B. Fractional Charge of Residues. C. Kappa, κ, charge residue clustering.

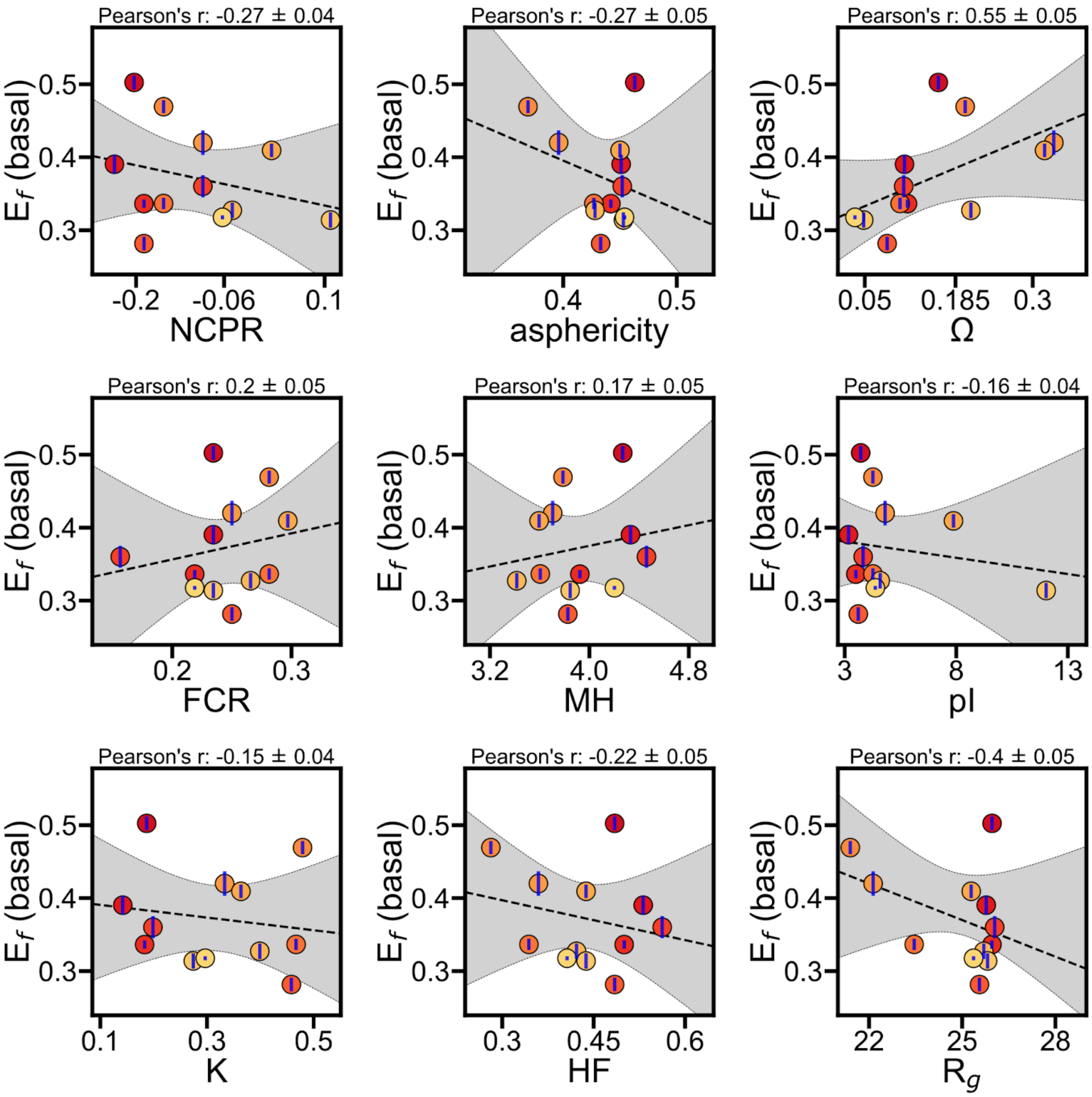


**Figure S3.** Computation scores predicted from cider and sparrow to experimental average $E_f$. Dashed line and shaded area represent a linear fit and 95% confidence intervals, respectively, and blue bars are standard deviations of $E_f$ experimental well medians shown in Fig 2C. The linear correlation is calculated for fraction of charged residues (FCR), net charge per residue (NCPR), charge clustering kappa (K), charge to proline omega (Ω), radius of gyration ($R_g$), isoelectric point (pI), mean hydropathy (MH), and hydrophobic fraction (HF).

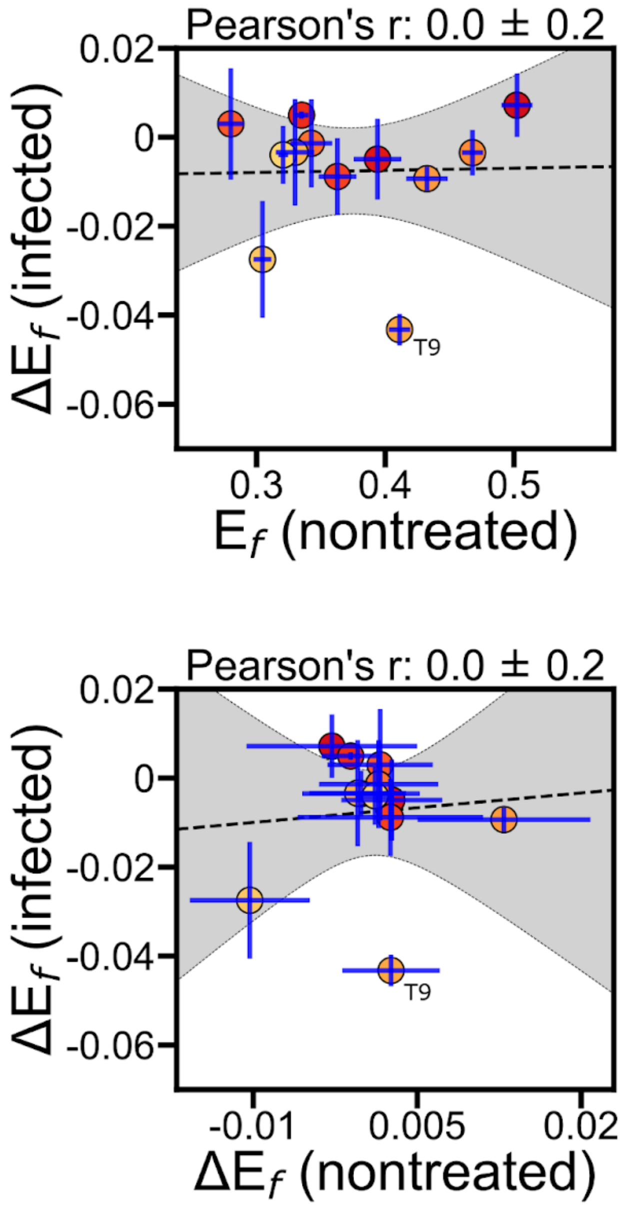


**Figure S4.** Experimental average values of basal nontreated $E_f$ at 48 hours and nontreated $\Delta E_f$ (Fig 3F). $\Delta E_f$ mean is calculated by averaging the change in well median values at 48 HPI by the well median values at 12 HPI ($\Delta E_f = E_f$ (48 HPI) - $E_f$ (12 HPI). Dashed line and shaded area represent a linear fit and 95% confidence intervals, respectively, and blue bars are standard deviations of $\Delta E_f$ experimental well medians at 48 HPI shown in Fig 3F. Tile 9 is labeled as “T9.”

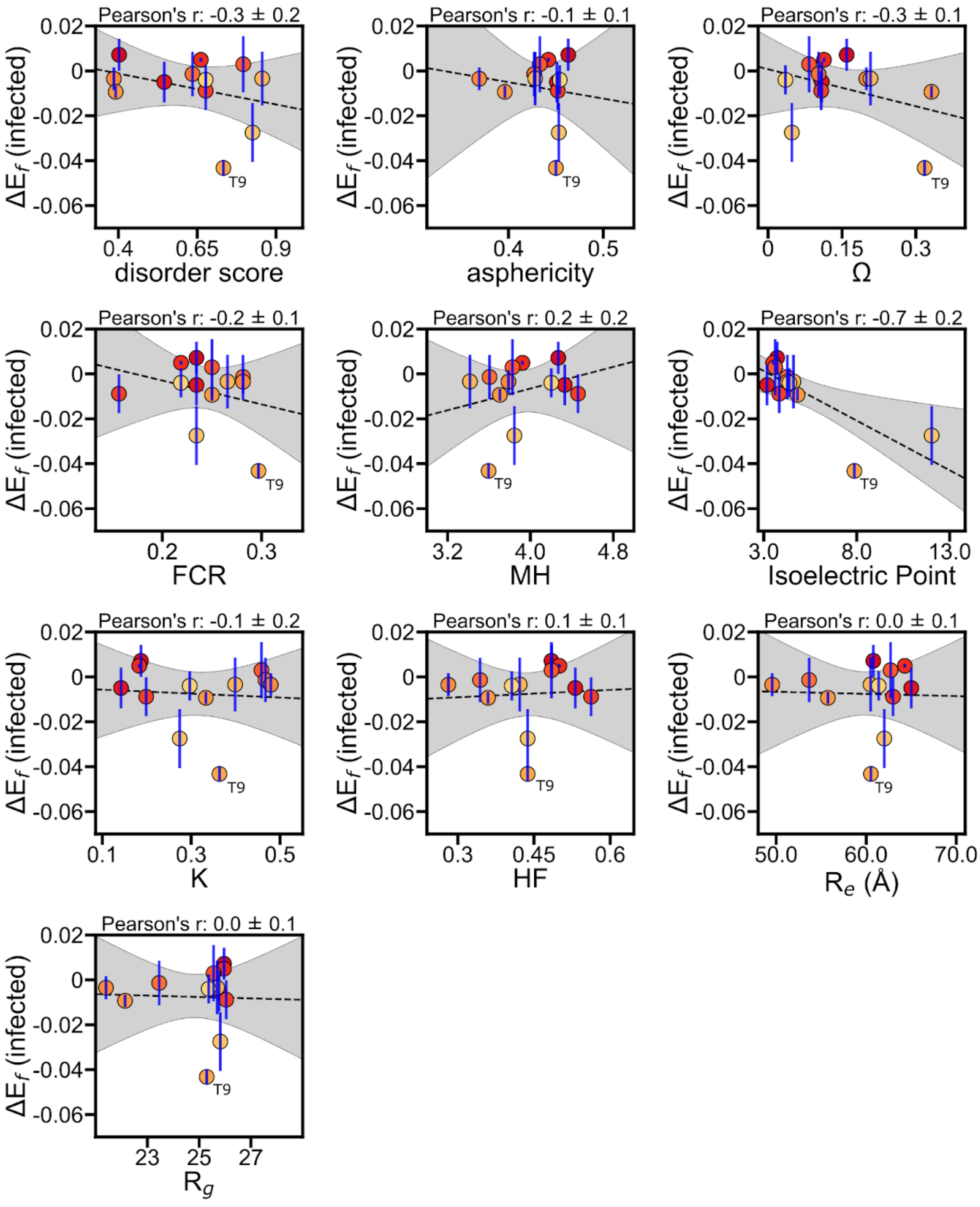


**Figure S5.** Computation scores predicted from cider and sparrow to experimental average $\Delta E_f$ in Fig. 3F. Dashed line and shaded area represent a linear fit and 95% confidence intervals, respectively, and blue bars are standard deviations of $\Delta E_f$ experimental well medians shown in Fig 3F. Computational comparisons are the same as in Fig. 1C and Fig. S3. Tile 9 is labeled "T9."

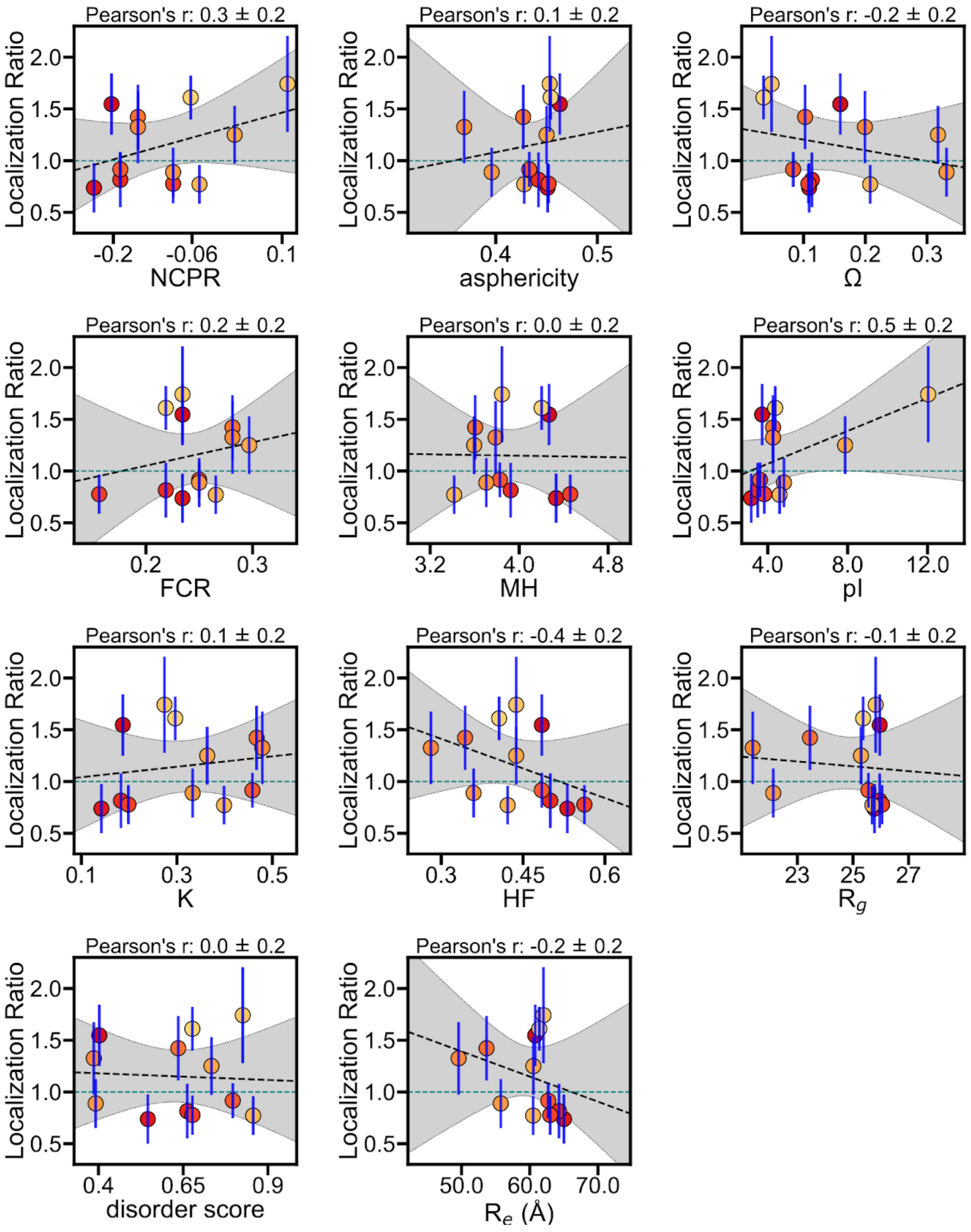


**Figure S6.** Computation scores predicted from cider and sparrow to median Localization Ratio in Fig. 4C. Dashed line and shaded area represent a linear fit and 95% confidence intervals, respectively, and blue bars are standard deviations of $\Delta E_f$ experimental well medians shown in Fig 4C. Computational comparisons are the same as in Fig. 1C and Fig. S3.

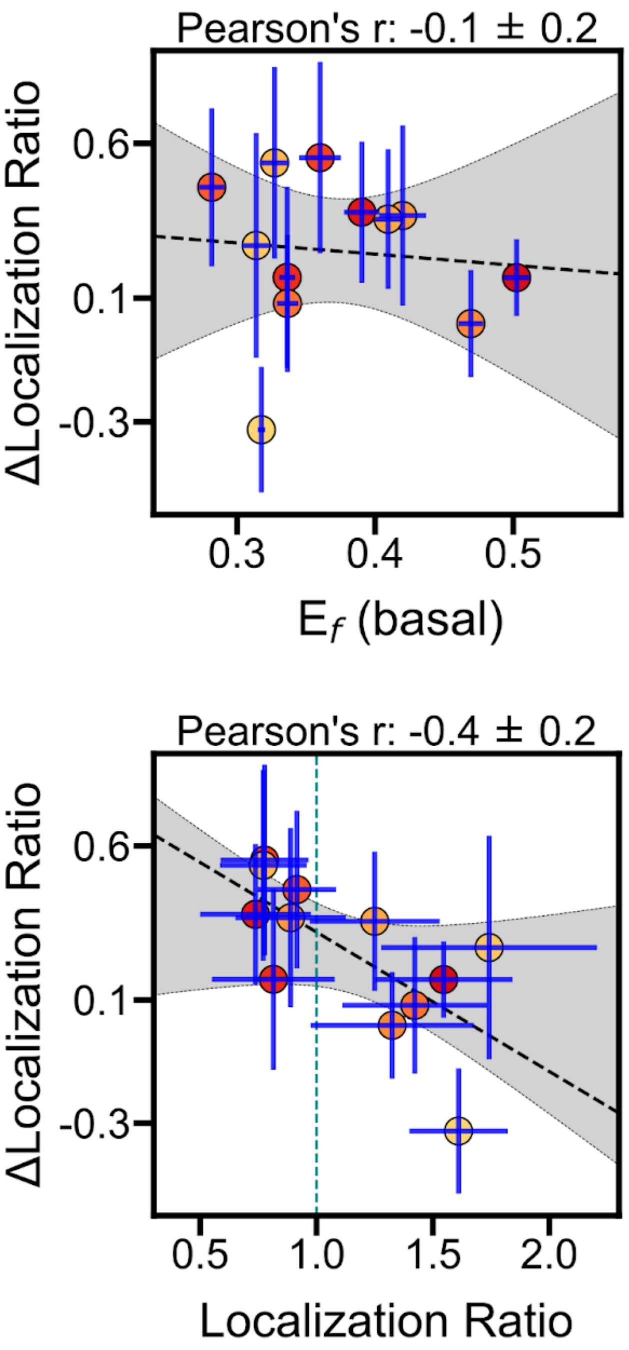


**Figure S7.** Comparative ratio of change in localization between population in Fig. 4F to experimental average basal $E_f$ population from Fig. 2C and localization in basal cell in Fig. 4C. ΔLocalization ratio is Localization Ratio (infected) - Localization Ratio (non-infected) at 48 HPI. Dashed line and shaded area represent a linear fit and 95% confidence intervals, respectively, and blue bars are standard deviations of experimental well medians shown in Fig. 2C, Fig. 4C, and Fig 4F.

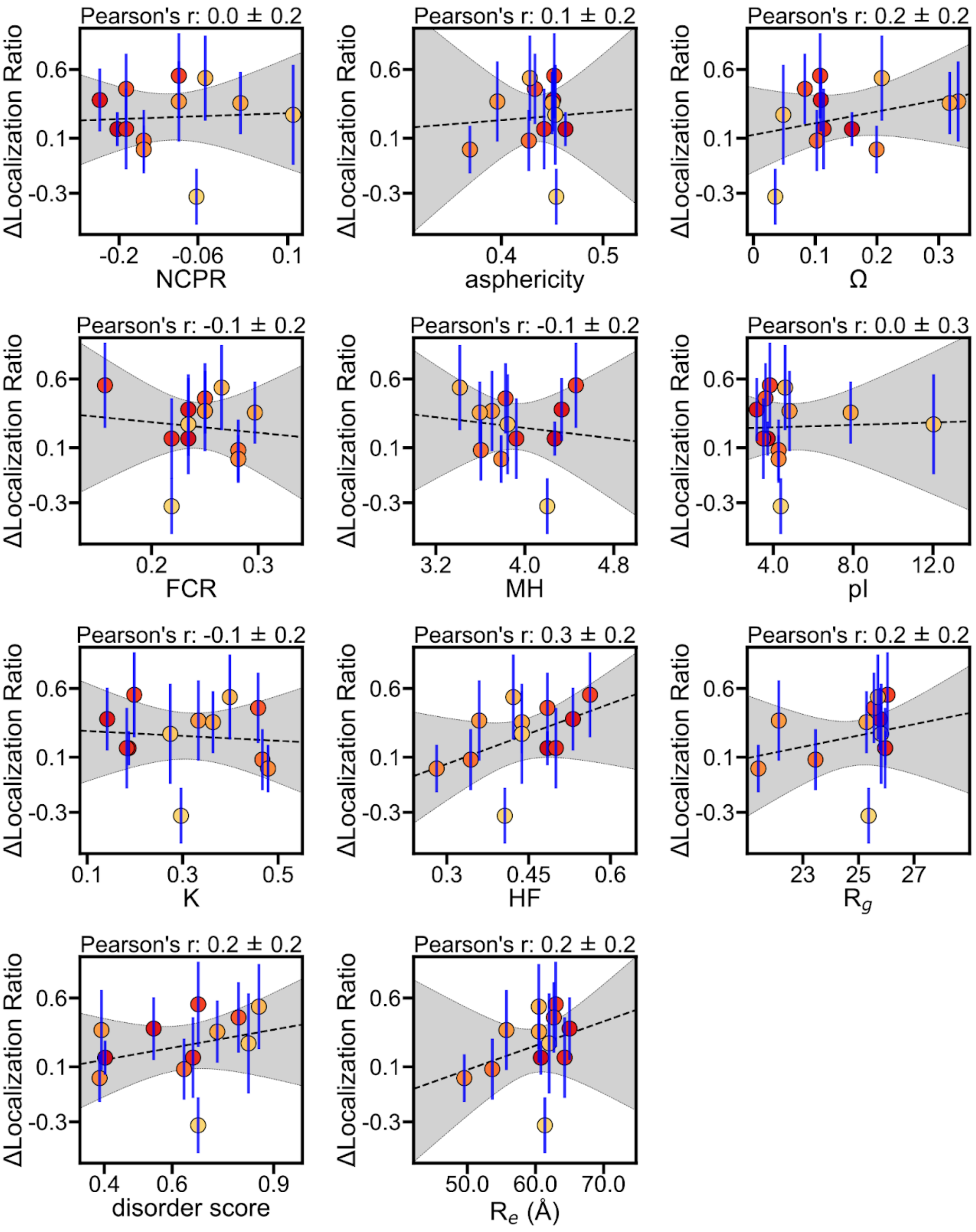


**Figure S8**. Computation Scores predicted from cider and sparrow to experimental average ΔLocalization Ratio based on median scores in Fig. 4F infected compared to noninfected. Dashed line and shaded area represent a linear fit and 95% confidence intervals, respectively, and blue bars are standard deviations of ΔLocalization Ratio experimental well medians shown in Fig 4F. Computational comparisons are the same as in Fig. 1C and Fig. S3.

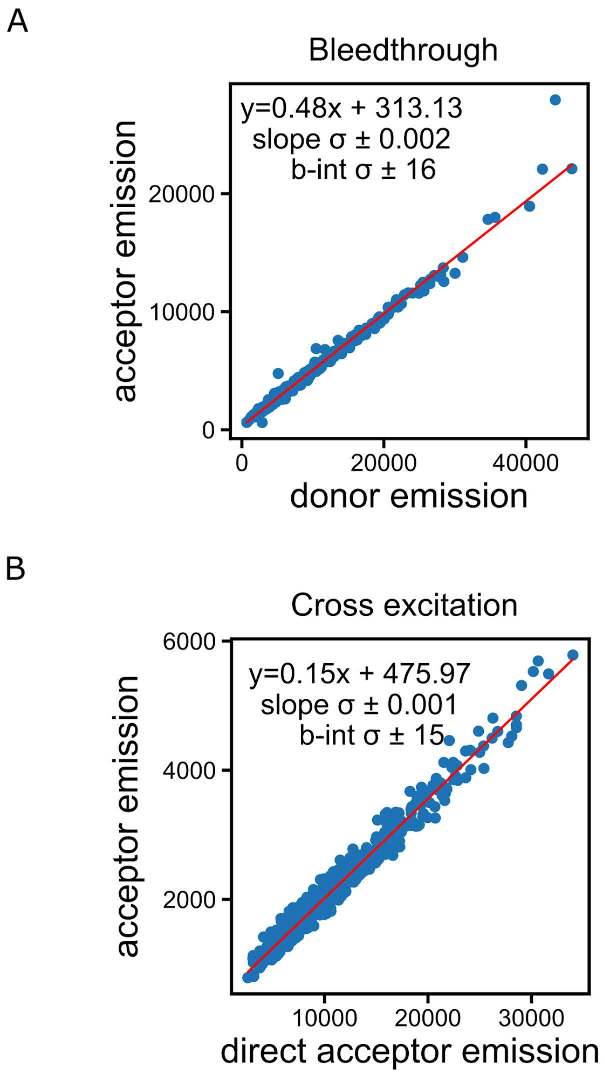


Fig S9. A. Bleedthrough and B. cross excitation correction values for FRET microscopy. Cell population (blue) and linear relationship (red). Slopes of lines (0.48, 0.15) are the used correction values.

| Motif Type | position | description | citation |
|---|---|---|---|
| LIG | 41-49 | pRB binding Groove | (4) |
| LIG | 53-91 | TAZ2 binding region | (5) |
| LIG | 112-117 | PxLxP SLiM | (6) |
| LIG | 122-126 | LxCxE | (7) |
| LIG | 154-174 | Zinc Binding Finger | (8, 9) |
| LIG | 279-283 | PxDLS SLiM | (10) |
| MOD | 89 | Serine Phosphorylation | (11) |
| MOD | 219 | Serine Phosphorylation | (12) |
| MOD | 231 | Serine Phosphorylation | (12) |
| TRG | 30-69 | N-terminal NLS | (13) |
| TRG | 70-80 | NES | (14) |
| TRG | 139-204 | CR3 NLS | (15) |
| TRG | 258-289 | Canonical C-terminal NLS | (15) |

**Table S1.** Experimentally validated Motifs of E1A. Motifs include cleavage sites (CLV), degrons (DEG), docking motifs (DOC), ligand binding motifs (LIG), post-translational modification sites (MOD), and targeting sequences (TRG).

| tile | start | end | sequence |
|---|---|---|---|
| 1 | 1 | 64 | MRHIICHGGVITEEMAASLLDQLIEEVLADNLPPPSHFEPPTLHELYDLDVTAPEDPNEEAVSQ |
| 2 | 21 | 84 | DQLIEEVLADNLPPPSHFEPPTLHELYDLDVTAPEDPNEEAVSQIFPDSVMLAVQEGIDLLTFP |
| 3 | 41 | 104 | PTLHELYDLDVTAPEDPNEEAVSQIFPDSVMLAVQEGIDLLTFPPAPGSPEPPHLSRQPEQPEQ |
| 4 | 61 | 124 | AVSQIFPDSVMLAVQEGIDLLTFPPAPGSPEPPHLSRQPEQPEQRALGPVSMPNLVPEVIDLTC |
| 5 | 81 | 144 | LTFPPAPGSPEPPHLSRQPEQPEQRALGPVSMPNLVPEVIDLTCHEAGFPPSDDEDEEGEEFVL |
| 6 | 101 | 164 | QPEQRALGPVSMPNLVPEVIDLTCHEAGFPPSDDEDEEGEEFVLDYVEHPGHGCRSCHYHRRNT |
| 7 | 121 | 184 | DLTCHEAGFPPSDDEDEEGEEFVLDYVEHPGHGCRSCHYHRRNTGDPDIMCSLCYMRTCGMFVY |
| 8 | 141 | 204 | EFVLDYVEHPGHGCRSCHYHRRNTGDPDIMCSLCYMRTCGMFVYSPVSEPEPEPEPEPEPARPT |
| 9 | 161 | 224 | RRNTGDPDIMCSLCYMRTCGMFVYSPVSEPEPEPEPEPEPARPTRRPKMAPAILRRPTSPVSRE |
| 10 | 181 | 244 | MFVYSPVSEPEPEPEPEPEPARPTRRPKMAPAILRRPTSPVSRECNSSTDSCDSGPSNTPPEIH |
| 11 | 201 | 264 | ARPTRRPKMAPAILRRPTSPVSRECNSSTDSCDSGPSNTPPEIHPVVPLCPIKPVAVRVGGRRQ |
| 12 | 221 | 284 | VSRECNSSTDSCDSGPSNTPPEIHPVVPLCPIKPVAVRVGGRRQAVECIEDLLNEPGQPLDLSC |

**Table S2.** E1A tiles. First and last residue, as well as full sequence are shown for each tile. Colorcode represents different residue types: positively charged (blue: R and K), negatively charged (red: E, D), aromatic (orange: F and Y), polar (green: H, G, S, Q, N, T), nonpolar (black: L, I, V, A, M), proline (purple), and cysteine (yellow)
